# Coral disease and ingestion: investigating the role of heterotrophy in the transmission of pathogenic *Vibrio* spp. using a sea anemone (*Exaiptasia pallida*) model system

**DOI:** 10.1101/2023.02.14.528589

**Authors:** William A. Norfolk, Carolina Melendez-Declet, Erin K. Lipp

## Abstract

Understanding disease transmission in corals can be complicated given the intracity of the holobiont and difficulties associated with *ex situ* coral cultivation. As a result, most of the established transmission pathways for coral disease are associated with perturbance (i.e., damage) rather than evasion of immune defenses. Here we investigate ingestion as a potential pathway for the transmission of coral pathogens that evades the mucus membrane. Using sea anemones (*Exaiptasia pallida*) and brine shrimp (*Artemia* sp.) to model coral feeding, we tracked the acquisition of the putative pathogens, *Vibrio alginolyticus, V. harveyi*, and *V. mediterranei* using GFP-tagged strains. *Vibrio* sp. were provided to anemones using three experimental exposures 1) direct water exposure alone, 2) water exposure in the presence of a food source (clean *Artemia*), and 3) through a “spiked” food source (*Vibrio*-colonized *Artemia*) created by exposing *Artemia* cultures to GFP-*Vibrio* via the ambient water overnight. Following a 3 h feeding/exposure duration, the level of acquired *GFP-Vibrio* was quantified from anemone tissue homogenate. Ingestion of spiked *Artemia* resulted in a significantly greater burden of GFP-*Vibrio* equating to an 829.7-fold, 3,108.2-fold, and 435.0-fold increase in CFU mL^−1^ when compared to water exposed trials and a 206.8-fold, 62.2-fold, and 27.3-fold increase in CFU mL^−1^ compared to water exposed with food trials for *V. alginolyticus, V. harveyi*, and *V. mediterranei*, respectively. These data suggest that ingestion can facilitate delivery of an elevated dose of pathogenic bacteria in cnidarians and may describe an important portal of entry for pathogens in the absence of perturbing conditions.

**Importance:** The front line of pathogen defense in corals is the mucus membrane. This membrane coats the surface body wall creating a semi-impermeable layer that inhibits pathogen entry from the ambient water both physically and biologically through mutualistic antagonism from resident mucus microbes. To date, much of the coral disease transmission research has been focused on mechanisms associated with perturbance of this membrane such as direct contact, vector lesions (predation/biting), and waterborne exposure through preexisting lesions. The present research describes a transmission pathway that evades the defenses provided by this membrane allowing unencumbered entry of bacteria as in association with food. This pathway may explain an important portal of entry for emergence of idiopathic infections in otherwise healthy corals and can be used to improve management practices for coral conservation.

## Introduction

In recent years, coral reefs have experienced unprecedented decline with regular mass mortality events occurring annually across the globe (Eddy et al., 2021). As ecosystem engineers, hermatypic corals produce the foundation of reef habitats by creating the critical three-dimensional structure that defines the reefscape (Wild et al., 2011). The loss of key coral species causes a decline in habitat complexity leading to a subsequent loss of biodiversity and reef ecosystem services (e.g., coastal protection, fisheries stability, and ecotourism) (Jones et al., 2004; Pratchett et al., 2018; Hoegh-Guldberg et al., 2019; Eddy et al., 2021). While coral decline can be attributed to many factors including global climate change, pollution, eutrophication, anthropogenic development, and overfishing, coral disease remains one of the most prominent causes of regional mortality events worldwide (Harvell et al., 1999; Green & Bruckner, 2000; Porter et al., 2001; Harvell et al., 2007; Montilla et al., 2019).

Understanding disease transmission, or how a pathogen spreads between individuals in a susceptible population, is a critical component for the management of infectious disease. A mechanistic understanding of the processes related to pathogen movement from reservoirs, through the environment, and into a susceptible host can provide insight for the prediction of disease outbreaks. Prior investigations of coral disease transmission have demonstrated the importance of direct contact, vector transmission, and waterborne transmission via preexisting lesions (reviewed by Shore & Cadwell, 2019). However, few studies have directly investigated the mechanisms of waterborne transmission, or ambient transmission via exposure in the water column, in uninjured healthy corals. Direct acquisition of pathogenic bacteria from the water column is impeded by the mucus membrane, which creates a semi-impermeable physical and biological barrier surrounding the coral tissue and by ciliary flows that create microscale water currents reducing the efficacy of pathogen chemotaxis (Rosenberg et al., 2007; Shapiro et al., 2014; Thompson et al., 2014). Thus, in the absence of injury where these systems are degraded, pathogens must overcome these defenses or utilize alternate portals of entry to establish infection.

Two recent studies have suggested that direct bacterial ingestion or ingestion of zooplankton may play an important role in the transmission of coral disease. Certner et al. (2017) demonstrated that white-band disease (WBD) transmission can be facilitated through zooplankton ingestion following incubation in tissue homogenate from diseased corals. In a similar vein, Gavish et al. (2021) utilized a microscale visualization system to observe colonization of *Pocillopora damicornis* by *Vibrio coralliilyticus* from ambient sea water, suggesting that ingestion may be a primary route of entry for the pathogen. Corals support their carbon and nutrient needs through the mutualistic relationship with their algal symbionts and through direct feeding. Heterotrophy provides up to 35% of a healthy coral’s daily metabolic needs and up to 100% in bleached corals, largely by nighttime feeding on zooplankton (Houlbrèque & Ferrier-Pagès, 2009; Ferrier-Pagès et al., 2010). While Gavish et al. (2021) demonstrates the viability of pathogen acquisition via direct ingestion of bacteria, preferential grazing of zooplankton, which are known to be colonized by bacteria (and *Vibrio* in particular [Erken et al., 2015]), may represent an important exploitable pathway for pathogenic microbes to gain entry to a coral host. We hypothesize that pathogen-colonized zooplankton may serve as a foodborne vector for disease transmission in uninjured corals.

*Vibrio* spp. are ubiquitous aquatic bacteria frequently identified as the causative or putative agents of coral disease (Table 1) (Munn, 2015). As indigenous microorganisms, or bacteria that exist naturally as a part of the ambient microbial community, *Vibrio* exhibit complex interspecies interactions that allow them to inhabit a broad range of ecological niches in the environment (Takemura et al., 2014). Of particular note is the association between *Vibrio* spp. and chitinous zooplankton (Takemura et al., 2014; Erken et al., 2015). Prior studies of *Vibrio* populations frequently associate total *Vibrio* and/or specific *Vibrio* spp. with plankton presence (Kaneko & Colwell, 1977; Heidelberg et al., 2002; Thompson et al., 2004; Turner et al., 2009; Magny et al., 2011; Martinez-Urtaza et al., 2011; Main et al., 2015). This association has been suggested to facilitate bacterial dispersal (Grossart et al., 2010; Erken et al., 2015), reduce bacterivore predation (Matz et al., 2005; Liang et al., 2019), and/or enable the utilization of chitin as a substrate (Hunt et al., 2008; Pruzzo et al., 2008; Erken et al., 2015).

**Table 1:** Published occurrences of *Vibrio* spp. as causative or putative agents of coral disease. Data organized by species. Recognized synonyms listed below. Updated from Kemp et al. (2018).

| *Vibrio* spp. | Disease Type | Disease Name or Description | Affected Host | Citation(s) |
| --- | --- | --- | --- | --- |
| *Vibrio coralliilyticus* | White disease<sup>a</sup> | Bacterial bleaching disease | *Pocillopora damicornis* and *Oculina patagonica* | (Ben-Haim et al., 2003)<br>(Rubio-Portillo et al., 2014) |
|  |  | *Montipora* white syndrome<sup>b</sup> | *Montipora capitata* | (Ushijima et al., 2014)<br>(Shore-Maggio et al., 2018) |
|  |  | Indo-Pacific white syndrome<sup>b</sup> | *Acropora cytherea*, *Montipora aequituberculata*, and *Pachyseris speciosa*. | (Sussman et al., 2008) |
| *Vibrio mediterranei* (*Vibrio shiloi*) | White disease<sup>a</sup> | Bacterial bleaching disease | *Oculina patagonica* | (Kushmaro et al., 2001)<br>(Thompson et al., 2001) |
| *Vibrio harveyi* (*Vibrio charchariae*) | White disease<sup>a</sup> | White band disease | *Acropora cervicornis* | (Ritchie & Smith, 1998)<br>(Gil-Agudelo et al., 2006) |
|  |  | White syndrome<sup>b</sup> | *Pocillopora damicornis* and *Acropora* spp. | (Marco Luna et al., 2010) |
|  | Yellow disease | Yellow band disease<sup>c</sup> | *Orbicella faveolata* | (Cervino et al., 2008) |
| *Vibrio alginolyticus* | White disease<sup>a</sup> | *Porites andrewsi* white syndrome | *Porites andrewsi* | (Zhenyu et al., 2013) |
|  | Yellow disease | Yellow band disease<sup>c</sup> | *Orbicella faveolata* | (Cervino et al., 2008) |
| *Vibrio natriegens* | White disease<sup>a</sup> | *Porites* ulcerative white spot disease | *Porites cylindrica* | (Arboleda & Reichardt, 2010) |
| *Vibrio owensii* | White disease<sup>a</sup> | *Montipora* White Syndrome<sup>b</sup> | *Montipora capitata* | (Ushijima et al., 2012)<br>(Shore-Maggio et al., 2018) |
| *Vibrio parahaemolyticus* | White disease<sup>a</sup> | *Porites* ulcerative white spot disease | *Porites cylindrica* | (Arboleda & Reichardt, 2010) |
| *Vibrio rotiferianus* | Yellow disease | Yellow band disease<sup>c</sup> | *Orbicella faveolata* | (Cervino et al., 2008) |
| *Vibrio tubiashii* | White disease<sup>a</sup> | White syndrome<sup>b</sup> | *Acropora muricata* | (Sweet & Bythell, 2015) |
| *Vibrio proteolyticus* | Yellow disease | Yellow band disease<sup>c</sup> | *Orbicella faveolata* | (Cervino et al., 2008) |
| Unspecified *Vibrio* spp. | Black disease | Black band disease<sup>b</sup> | *Favia* spp. | (Arotsker et al., 2009) |
|  | White disease<sup>a</sup> | Stony coral tissue loss disease<sup>b</sup> | *Montastraea cavernosa*,<br>*Orbicella faveolata*,<br>*Diploria labyrinthiformis*,<br>and *Dichocoenia stokesii* | (Meyer et al., 2019) |

<sup>a</sup>Described by different authors under the names white, syndrome, pox, and/or band disease.
Disease signs are manifestations of coral tissue loss and/or zooxanthellae loss or bleaching.
<sup>b</sup>Associated as a part of a bacterial consortium suspected to contain non-*Vibrio* species.
<sup>c</sup>Associated as a part of a bacterial consortium suspected to contain multiple *Vibrio* spp.

Research investigating cholera transmission in humans has demonstrated that *V. cholerae* cells colonize the exoskeletons of copepods where their concentration can increase to an excess of 10^4^ cells copepod^−1^ (Huq et al., 1983; Tamplin et al., 1990; Colwell, 1996; Rawlings et al., 2007; Magny et al., 2011). Subsequent ingestion of colonized copepods can increase the probability of ingesting a potentially pathogenic dose of the bacterium facilitating the onset of disease (Huq et al., 1996; Nelson et al., 2009). Furthermore, pre-filtration of surface water sources utilized for drinking with simple fabric mesh can reduce the occurrence of *V. cholerae* infections due to the reduction of colonized zooplankton (Huq et al., 1996; Colwell et al., 2003). While this *Vibrio*-zooplankton transmission pathway has been well established for *V. cholerae*, little research has been devoted to investigating the importance of these interactions for non-cholera *Vibrio* infections.

The work presented here investigates the viability of an ingestion-based transmission pathway for the acquisition of potentially pathogenic *Vibrio* spp. in corals. To alleviate difficulties of *ex situ* coral cultivation, a model system was employed utilizing sea anemones (*Exaiptasia pallida*) and brine shrimp (*Artemia* sp.) to mimic natural coral feeding. Prior research has demonstrated the utility of sea anemones in the genus *Exaiptasia* (formally *Aiptasia*, see Grajales & Rodriguez, 2014 for reclassification) as lab-friendly surrogates for coral experimentation (Belda-Baillie et al., 2001; Weis et al., 2008; Sunagawa et al., 2009; Hardefeldt & Reichelt-Brushett, 2015). Structurally, *Exaiptasia* spp. resemble large non-colonial coral polyps and feed both heterotrophically on zooplankton and autotrophically though the use of their algal symbionts (zooxanthellae) (Grajales & Rodriguez, 2014). Using this model system, we traced the acquisition of the putative coral pathogens *V. alginolyticus, V. mediterranei*, and *V. harveyi*.

## Results

### *Artemia* Colonization by *Vibrio*

Colonization experiments first assessed the ability of *Vibrio* spp. to attach to/associate with *Artemia*. Substantial colonization of *Artemia* gastrointestinal (GI) tracts was observed for all tested vibrios following overnight (18 h) exposure via ambient water. Total colonization for each *Vibrio* spp. exposure (~250 *Artemia*) was 4.90 × 10^6^, 1.47 × 10^6^, 7.59 × 10^6^ CFU per ~250 individuals for *V. alginolyticus, V. harveyi*, and *V. mediterranei*, respectively. These levels equate to a mean acquisition of 4.32%, 2.14%, and 50.21% of the initial exposure dose for *V. alginolyticus, V. harveyi*, and *V. mediterranei*, respectively (Figure S1). Epifluorescence microscopy showed GFP tagged cells were concentrated throughout the length of *Artemia* GI tracts in association with ingested material and feces (Figure 1). GFP cells were also observed in association with *Artemia* feces following defecation. Low exoskeletal association was observed in all experimental trials, though minor attachment and/or entanglement was noted in association with *Artemia* appendages (Figure 1). GI association was consistent across naupliiar sizes excluding the smallest, most recently hatched individuals (Figure S2), which showed little to no *GFP-Vibrio* accumulation. Visual patterns of GI association did not differ between *Vibrio* species. No distinctive behavioral changes or swimming impairment was observed in colonized *Artemia* throughout the duration of exposure (up to 24 h).

**Figure 1:**
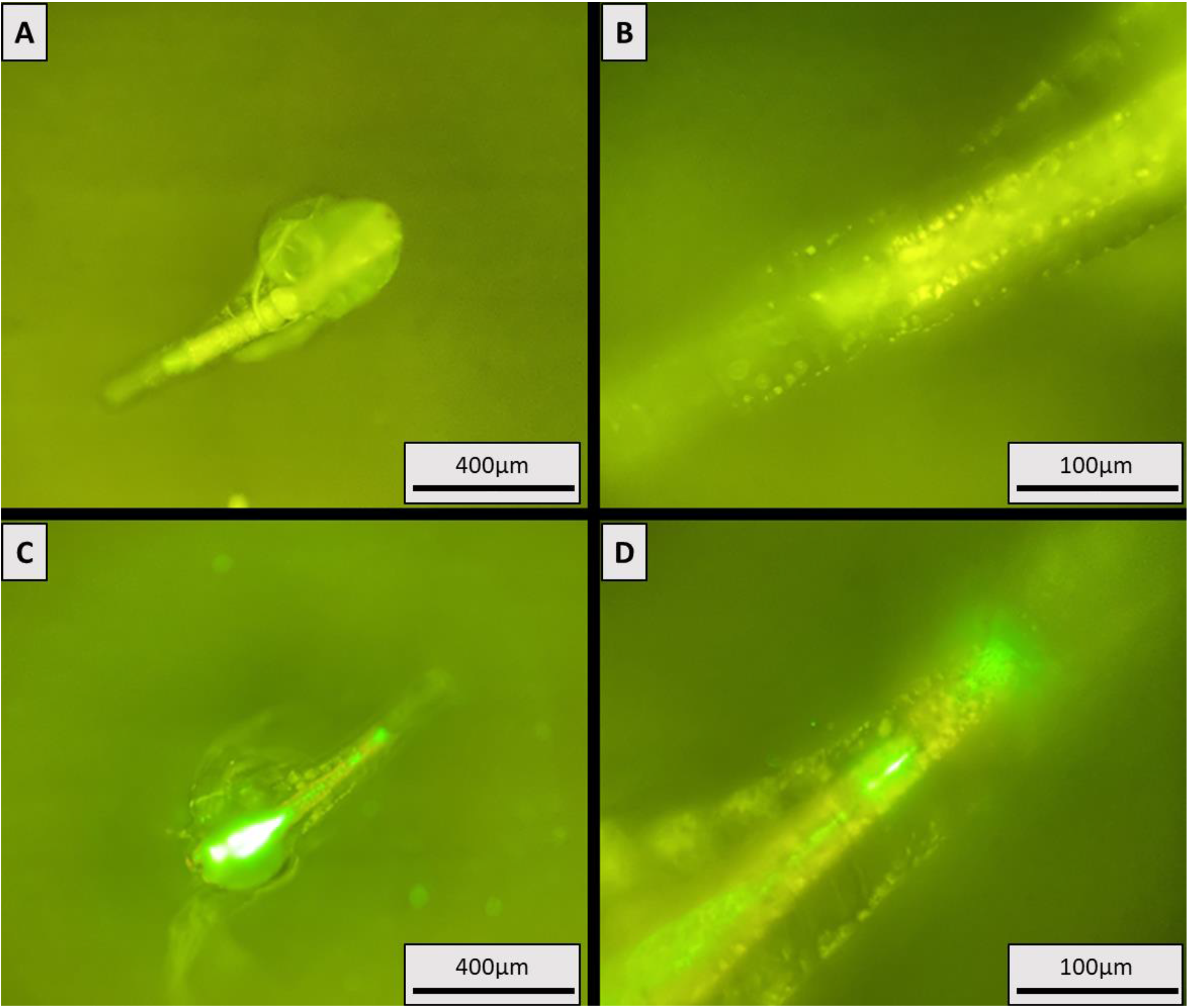
GFP *V. alginolyticus* colonization of *Artemia*. Cultures inoculated with ~1.13 × 10^8^ CFU. Photos taken after 18 h of exposure. (A) Unexposed *Artemia* at 100X magnification. (B) Unexposed *Artemia* posterior at 400X magnification. (C) Exposed *Artemia* at 100X magnification. (D) Exposed *Artemia* posterior at 400X magnification. Bright green fluorescence indicates GFP *V. alginolyticus* presence. *Artemia* tissue appears yellow green.

### Uptake of *Vibrio* by *E. pallida*

Anemone feeding studies evaluated the efficacy of an ingestion-based transmission pathway by confirming consumption of GFP-*Vibrio-*colonized *Artemia* and quantification of the acquired GFP-*Vibrio* dose. Gross observations of feeding demonstrate that *E. pallida* readily ingested *Vibrio*-colonized *Artemia*, responding rapidly with predatory tentacle behavior when *Artemia* were introduced into the microcosm water (Figure S3). No differences in anemone feeding behavior (i.e., tentacle response) were observed for exposures using spiked and non-spiked *Artemia*.

Assessment of the acquired dose compared four major feeding/exposure treatments: 1) water exposed not fed, where GFP-*Vibrio* were inoculated into the microcosm water and no *Artemia* were added, 2) water exposed control fed, where GFP-*Vibrio* were inoculated into the microcosm water and anemones were fed with clean (non-spiked) *Artemia*, 3) spiked fed, where no GFP-*Vibrio* were inoculated into the microcosm water and anemones were fed with *Vibrio*-colonized *Artemia*, and 4) control, where no GFP-*Vibrio* were inoculated into the microcosm water and anemones were fed clean *Artemia* (Figure 2). Significantly greater GFP-*Vibrio* levels were observed in *E. pallida* individuals exposed via spiked *Artemia* (spiked fed) compared to individuals exposed through the ambient water, regardless of the presence of *Artemia* (i.e., all other experimental conditions). Anemone homogenate from spiked fed trials showed a mean GFP-*Vibrio* concentration of 6.92 × 10^4^, 2.59 × 10^5^, and 1.67 × 10^5^ CFU mL^−1^ for *V. alginolyticus*, *V. harveyi*, and *V. mediterranei*, respectively. Conversely, water exposed anemones showed a mean concentration of 3.33 × 10^2^ and 8.33 × 10^1^ CFU mL^−1^ for *V. alginolyticus*, 4.10 × 10^3^ and 8.33 × 10^1^ CFU ml^−1^ for *V. harveyi*, and 5.90 × 10^3^ and 3.83 × 10^2^ CFU mL^−1^ for *V. mediterranei* for water exposed control fed (non-spiked) and water exposed not fed (no *Artemia*) treatments, respectively. These concentrations equate to a 206.8-fold, (p-value = 0.03), 62.2-fold (p-value = 0.013), and 27.3-fold (p-value = 0.013) increase in the GFP-*Vibrio* burden of spiked fed compared to water exposed control fed anemones and a 829.7-fold (p-value = 0.028), 3,108.2-fold (p-value = 0.026), and 435.0-fold (p-value = 0.030) increase in spiked fed compared to water exposed not fed anemones for *V. alginolyticus*, *V. harveyi*, and *V. mediterranei*, respectively (Figure 3). Between the two water exposures, fed (non-spiked *Artemia*) anemones showed a significantly greater burden of GFP *V. harveyi* (p-value = 0.026) and *V. mediterranei* (p-value = 0.030) compared to non-fed anemones but did not differ significantly for *V. alginolyticus* (p-value = 0.51). No GFP-*Vibrio* were recovered from anemones in the control group (no exposure) or from anemone wash water (carry-over control).

**Figure 2:**
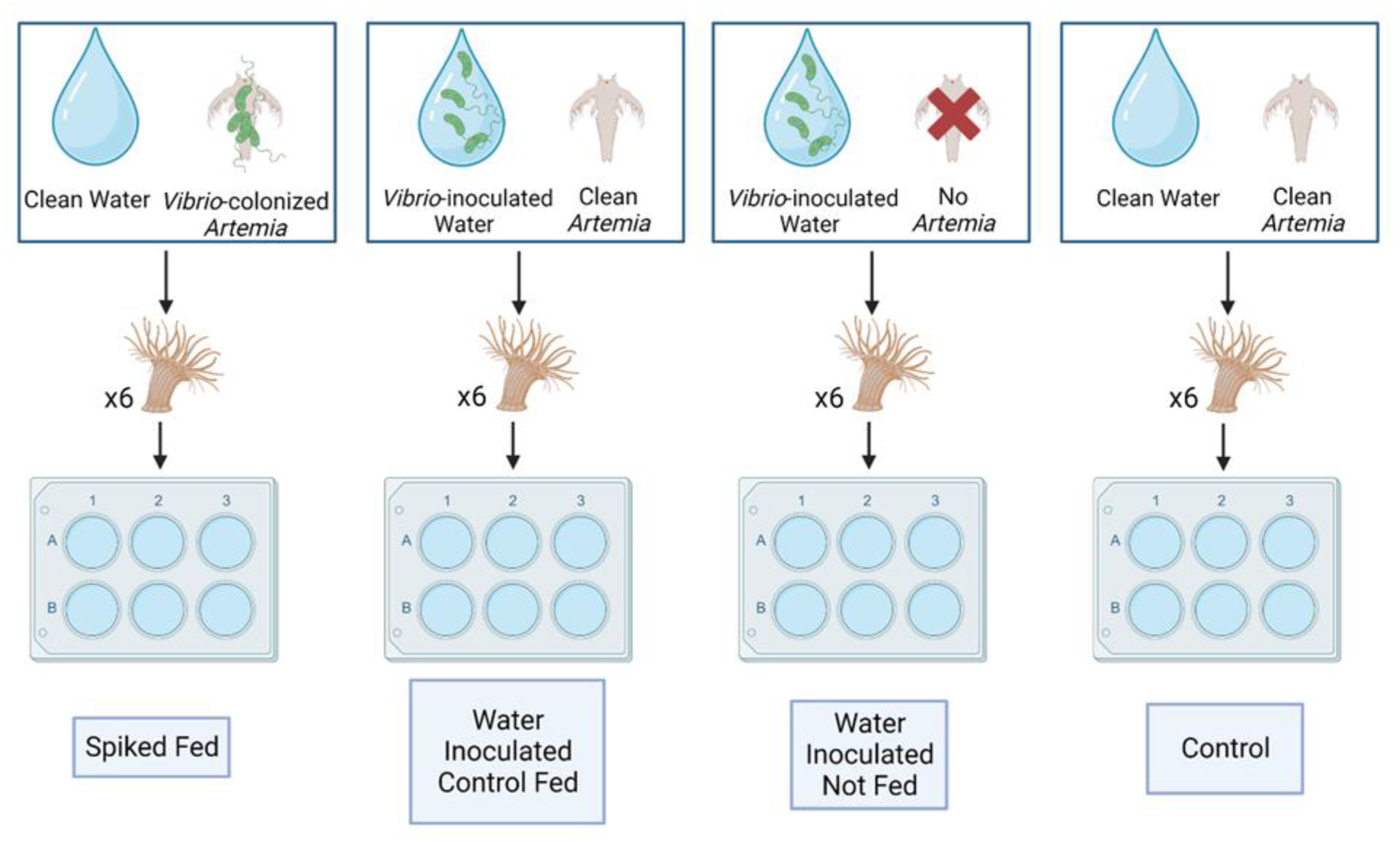
Feeding trial treatments used to expose *E. pallida* to GFP-*Vibrio*. *Artemia* administered at a concentration of ~1,000 individuals (when added). GFP-*Vibrio* administered at concentrations designated in Table 2.

**Figure 3:**
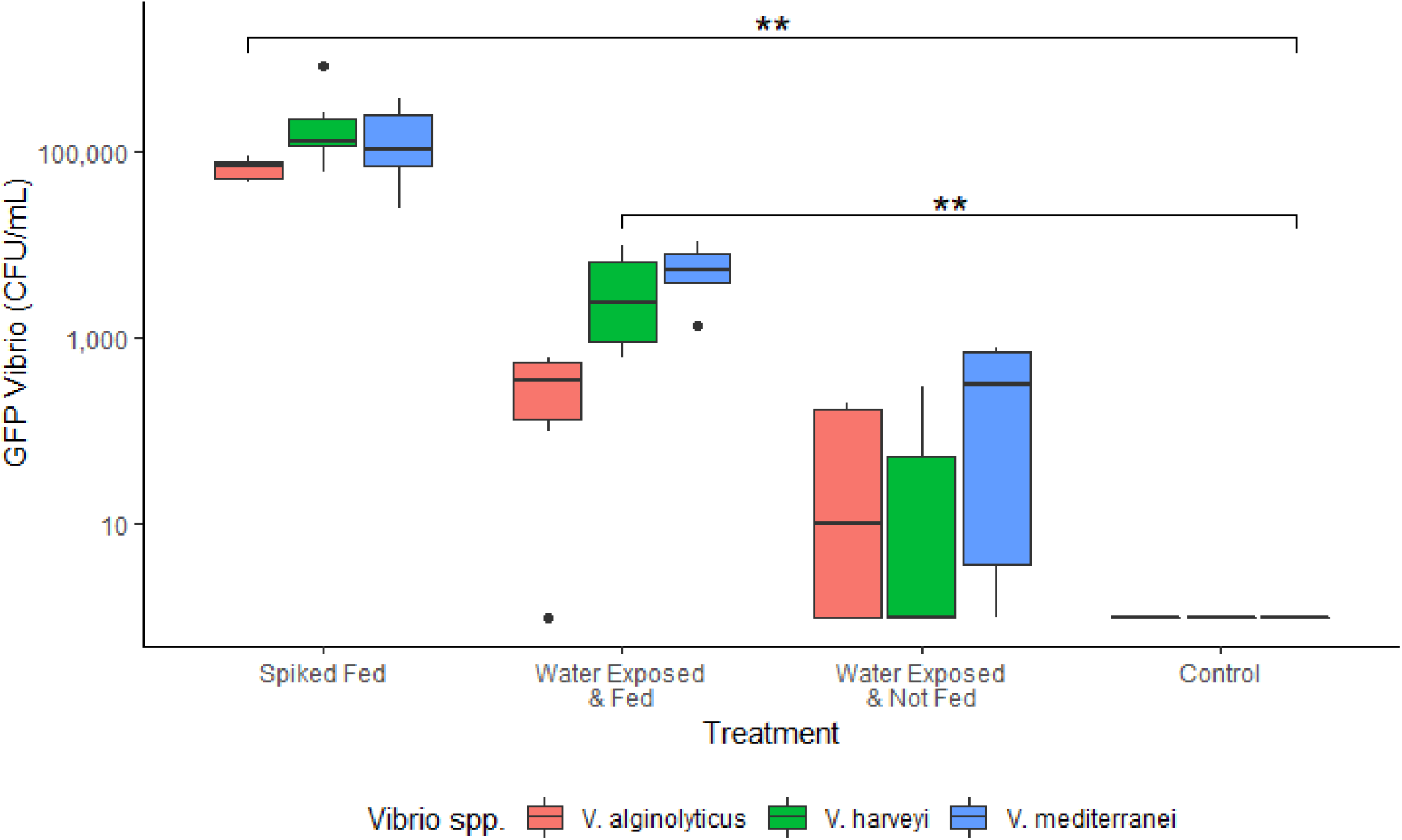
Recovered *GFP-Vibrio* spp. concentrations from anemone homogenate following completion of the controlled feeding study. Spiked fed anemones demonstrated a significantly greater *GFP-Vibrio* spp. burden compared to water exposed individuals. Spiked fed versus water exposed and fed resulted in p-values of 0.03, 0.013, and 0.013 and spiked fed versus water exposed not fed resulted in p-values of 0.0047, 0.0043, and 0.0049 for *V. alginolyticus, V. harveyi*, and *V. mediterranei*, respectively. N = 6 anemones for each exposure type and *Vibrio* spp.

## Discussion

The mucus membrane serves as the front line of defense against infection for coral species. This mucus coats the epithelia creating a semi-impermeable barrier between the coral tissue and ambient water (Cooney et al., 2002; Brown & Bythell, 2005; Rosenberg et al., 2007; Thompson et al., 2014). Within this mucus layer, a variety of mutualistic and commensal microorganisms are maintained. The totality of these microbes and the coral colony are collectively known as the holobiont (Rohwer et al., 2002). Research has suggested that the coral-associated microbial community can confer improved fitness to the holobiont through community shifts in response to environmental change (Reshef et al., 2006, Thompson et al., 2014), the production of antimicrobial compounds (Shnit-Orland & Kushmaro, 2009) and/or antagonistic competition with potential pathogens (Rohwer et al., 2002; Ritchie, 2006; Teplitski & Ritchie, 2009). Together, the physical mucus barrier combined with the biological protection of the microbial community pose a substantial challenge to the direct transmission of waterborne pathogens. To date, the majority of coral disease transmission research has focused on mechanisms of pathogen spread associated with perturbance of this mucus membrane such as direct contact, vector-mediated (i.e., biting), and indirect transmission via preexisting lesions (Shore & Cadwell, 2019). While these studies provide important insight into the ecology of coral diseases, these transmission mechanisms are dependent on opportunistic occurrences related to host proximity and preexisting or active damage and there is substantial need to investigate transmission mechanisms related to disease emergence in uninjured corals.

Despite the presence of zooxanthellae, heterotrophic feeding is a critical component of coral nutrition, accounting for up to 35% of the daily metabolic needs of some coral species (Houlbrèque & Ferrier-Pagès, 2009; Ferrier-Pagès et al., 2010). Corals preferentially feed on small zooplankton thus, we investigated the ability of pathogenic *Vibrio* spp. to be transmitted to a cnidarian host via ingestion following colonization of a zooplankton vector. Using sea anemones (*E. pallida*) and brine shrimp (*Artemia* spp.) to model coral feeding, we demonstrate that ingestion of *Vibrio*-spiked brine shrimp results in a significantly higher bacterial burden in recipient anemones compared to ambient water exposures, both with and without food sources (i.e., *Artemia*). These data suggest that ingestion could play a role in the transmission of certain coral pathogens. Furthermore, this mode of transmission bypasses the natural defense mechanisms of corals provided by their mucus membrane (Rohwer et al., 2002; Shnit-Orland & Kushmaro, 2009) which may describe an important portal of entry related to pathogenic infection of uninjured corals.

Acting as our model zooplankton, *Artemia* were readily colonized by all tested *Vibrio* spp. following direct waterborne exposure, similar to previous studies in *V. cholerae* (Huq et al., 1983). However, the preferential colonization of the GI tract noted here differed from previously described observations where colonization was predominately observed on zooplankton exoskeletons (Huq et al., 1983; Tamplin et al., 1990; Colwell, 1996). We hypothesize that this difference may be due to the fact that the present research was conducted *ex situ* where certain environmental determinants of zooplankton colonization (i.e., substrate limitation) may not be present and/or as impactful (Worden et al., 2005; Takemura et al., 2014, Liang et al., 2019). While some minor exoskeletal association was observed on *Artemia* appendages, we suspect that this may be the result of incidental entanglement rather than purposeful attachment. Due to the lack of strong external association, we postulate that the colonization of *Artemia* GI tracts is the result of active ingestion of *Vibrio* spp. by nauplii occurring over prolonged interaction (≥4 h of exposure). This hypothesis is further supported by the observation that the smallest most recently hatched *Artemia* (Figure S2) showed minimal GI colonization. At this stage of life, nauplii are nutritionally maintained through residual yolk protein and do not actively feed until they are larger (Warner et al., 1973; Sugumar & Munuswamy, 2006). The total *Artemia*-acquired dose remained relatively consistent for all three *Vibrio* spp. at ~10^6^ CFU per ~250 individuals. These data suggest that *Artemia* have a threshold for the maximum concentration of *Vibrio* spp. they can harbor via GI colonization.

Feeding experiments demonstrate that spiked fed anemones acquire a significantly greater GFP-*Vibrio* burden compared to water exposed individuals regardless of the presence of food. This pattern was observed across all three *Vibrio* spp. suggesting that ingestion of *Vibrio*-colonized zooplankton can facilitate delivery of an elevated dose of these bacteria, broadly. The higher *Vibrio* levels are likely the result of bioaccumulation of these bacteria within *Artemia* facilitating acquisition of a highly concentrated dose through targeted feeding. This is consistent with prior observations of *V. cholerae* carriage by copepods where ingestion of a small number of individuals may facilitate receipt of a potentially pathogenic dose (≤10^3^ cells) (Huq et al., 1983; Colwell, 1996). While low compared to spiked fed individuals, water exposed anemones did result in some uptake of GFP-*Vibrio* with higher levels acquired in the presence of food (‘clean’ *Artemia*) than without. This observation is consistent with the findings of Gavish et al. (2021) and suggests that even in the absence of *Vibrio*-colonization of food sources, active feeding and ingestion may contribute to the acquisition of *Vibrio* spp. cells from the surrounding water.

At ambient levels, *Vibrio* spp. typically range from 10^1^ to 10^3^ CFU mL^−1^ (Urakawa and Rivera, 2006) with location-specific differences in community composition driven largely by temperature and salinity (Turner et al., 2009; Takemura et al., 2014). However, *Vibrio* populations are known to be dynamic, fluctuating on a “boom-bust” cycle of growth and reduction in association with ephemeral pulses of limiting nutrients (Westrich et al., 2016; Westrich et al., 2018; Borchardt et al., 2020). During bloom events, total *Vibrio* can increase dramatically rising to levels 5 to 30 times greater than the typical background concentration of coastal waters (Westrich et al., 2016). Prior research has shown that seasonal increases in *Vibrio* abundance facilitate increases in both free-living and zooplankton-associated abundance (Carli et al., 1993). Thus, blooms numbers could potentially promote zooplankton colonization and enhance the likelihood of transmission via ingestion during these events. While further studies on species-specific colonization rate, transmitted dose, and uptake *in situ* are needed to assess the potential importance in coral disease, we postulate that these mechanisms provide an ecological basis for foodborne transmission of certain coral pathogens.

While the scope of this research is targeted at understanding coral disease, the results of this study have broader implications for the spread of vibriosis. *Vibrio* spp. have been implicated as the causative or putative pathogens in numerous diseases of marine organisms, most notably important aquaculture species such as Pacific White Shrimp (*Litopenaeus vannamei*), Tiger Prawn (*Penaeus monodon*), Atlantic Salmon (*Salmo salar*), and Gilt-Head Sea Bream (*Sparus aurata*) (Karunasagar et al., 1994; Press & Lillehaug, 1995; Balebona et al., 1998; Zhou et al., 2012). Zooplankton serve as the base of the marine/estuarine food web thus, there is potential for ingestion to play a role in the acquisition of these and similar pathogens. This hypothesis is supported by the work of Goulden et al. (2012) who utilized a similar GFP tracking system to demonstrate that *Panulirus ornatus* (ornate spiny lobster) mortality can be facilitated by ingestion of *V. owensii*-colonized *Artemia* in aquaculture settings. Furthermore, the non-discriminant acquisition of all three *Vibrio* spp. in the present study suggests that this pathway may be broadly viable within the Vibrionaceae and warrants continued investigation of the role of ingestion in the spread of other pathogenic vibrios.

## Conclusion

Understanding coral disease transmission is critical to the conservation of reef habitats. The present study describes a mechanistic pathway for the acquisition of coral pathogens via zooplankton ingestion using a sea anemone (*E. pallida*) and brine shrimp (*Artemia*) model system to represent coral heterotrophy. The results of this research demonstrate that ingestion of *Vibrio*-colonized *Artemia* can facilitate receipt of a significantly elevated *Vibrio* dose when compared to exposure via the water column suggesting that heterotrophy may represent an important transmission pathway for certain coral pathogens. Characterization of this pathway illustrates a means by which pathogenic bacteria may bypass the natural immune defenses of corals conferred by their mucus membranes allowing for unencumbered acquisition of a pathogenic dose. This mechanism may help to explain a potential source of idiopathic infections that arise in otherwise healthy unperturbed corals.

## Methods

### Experimental Vibrio Strains

Experimental *Vibrio* strains were obtained from our culture collection (E.K. Lipp, University of Georgia) and consisted of the known coral pathogens *V. alginolyticus, V. mediterranei*, and the putative coral pathogen *V. harveyi* (see Table S1 for strain information). All strains were maintained at −80°C in a 1:1 mixture of 40% glycerol (20% final concentration) and lysogeny broth (LB, Sigma Aldrich, Miller formulation) amended to 3% w/v NaCl (termed LBS 3%). To revive from storage, strains were inoculated into 4 mL LBS 3% and incubated at 30°C with 100 rpm shaking agitation (New Brunswick Scientific, C24 Incubator Shaker) for 18-24 h.

### Brine Shrimp Cultures and Maintenance

*Artemia* sp. were purchased as dehydrated cysts (Premium Grade Brine Shrimp Eggs: Brine Shrimp Direct Inc., Great Salt Lake Origin, USA). Dehydrated cysts (0.3 g) were revived in 300 mL sterile artificial sea water (35 practical salinity units [PSU] Instant Ocean^®^, termed ASW) incubated at room temperature under mild agitation from an aquarium bubbler (Whisper^®^ 20, Aquarium Air Pump). Cysts hatching occurred within 1-2 days of rehydration. *Artemia* were harvested at the nauplii stage, following 1-2 additional days of incubation, using a sterile serological pipette. Free swimming nauplii were collected from below the water surface to reduce collection of any discarded or unhatched cysts. Any *Artemia* cultures that appeared discolored (cloudy water), produced poorly swimming nauplii, or hatched insufficiently (<75% hatching, estimated visually) were discarded.

### Anemone Cultures and Maintenance

*E. pallida* anemones were purchased live (Carolina^®^ Biological Supply, #162865) and maintained in laboratory holding tanks. Holding tanks were constructed using a 6 L glass aquarium equipped with a constant-flow water filter (Aqueon^®^ QuietFlow Aquarium Power Filter 10), an in-water aquarium heater (Aqueon^®^ Pro Heater 50W), and a 445nm aquarium light (GloFish^®^ Blue, LED Aquarium Light). Holding aquaria were maintained under the conditions outlined in Tables S2 and S3. Prior to experimentation, all anemones were transferred to holding tanks and allowed to acclimate for a minimum of 2 weeks. Anemones were monitored daily, and any deceased individuals were removed. Long-term cultures (not used for experimentation) of *E. pallida* were kept with the experimental anemones to stabilize holding tank water chemistry. While in the holding tank, anemones were fed twice per week with 50 mL (~2,000 individuals) of decapsulated *Artemia*. Water changes (50% of tank volume) were preformed every two weeks and replaced volumetrically with fresh ASW. Intermittent tank cleaning was performed as needed using a scrub brush and/or a serological pipette to remove anemone debris and algal build-up following feeding.

### GFP Tagging

All *Vibrio* spp. used in this experiment were tagged with GFP to enable localization and quantification of the bacterium. Tagging was accomplished using the methods outlined in Norfolk & Lipp, (2022). In short, a tri-parental mating assay was used to transfer a *gfp*-containing plasmid to the target *Vibrio* sp. using bacterial conjugation. In this assay, two strains of *Escherichia coli*, the helper strain carrying the conjugative plasmid pEVS104 (*tra trb* Kn^r^) and the donor strain carrying the *gfp* plasmid pVSV102 (*gfp* Kn^r^), were combined in culture with the target *Vibrio* under mild kanamycin stress to promote transfer of the *gfp* plasmid. Fluorescence of all transconjugant (GFP-tagged) *Vibrio* spp. was confirmed using fluorescent microscopy (Olympus BX41 Fluorescence Microscope). Working stocks of transconjugant strains were maintained at room temperature in deep agar stabs containing LBS 3% amended with 300 μg mL^−1^ kanamycin to ensure retention of the plasmid. GFP strains were maintained at −80°C in a 1:1 mixture of 40% glycerol and LBS 3% broth amended with 300 μg mL^−1^ kanamycin for long-term storage.

### *Artemia* Colonization

*GFP-Vibrio* spp. were revived from −80°C storage in 4 mL of LBS 3% broth amended with 300 μg mL^−1^ kanamycin and incubated at 30 °C with 100 rpm of shaking agitation for 18-24 h. Following incubation, 1mL of the overnight culture was pelleted by centrifugation at ~4,000 × g, the supernatant was discarded, and replaced with 1 mL of sterile 1X phosphate buffered saline (PBS). This process was repeated three times to ensure adequate removal of residual kanamycin from the culture. Concurrently, *Artemia* cultures were grown as described above. Six mL of decapsulated nauplii (~250 individuals) were transferred to each well of a sterile six-well tissue culture plate (Cellstar^®^ 6-Well Suspension Culture Plate). Each well of the culture plate was inoculated with 50 μL of washed GFP *V. alginolyticus* (~1.13 × 10^8^ CFU), *V. harveyi* (~6.87 × 10^7^ CFU), or *V. mediterranei* (~1.51 × 10^7^ CFU). The *Artemia-Vibrio* mixture was covered and incubated at 28 °C under 50 rpm of shaking agitation for 18 h. Following incubation, the contents of each well was collected and filtered onto a 3.0 μm polycarbonate (PCTE) membrane (Sterlitech© 47mm, 3.0μm PCTE membranes) using vacuum filtration to capture the suspended *Artemia* while allowing any non-associated *Vibrio* cells to be discarded as flow through. The *Vibrio*-colonized *Artemia* were resuspended from the membrane by vortexing for 30 s in 6 mL of sterile ASW. Colonization or apparent attachment of GFP-labeled cells to *Artemia* nauplii was confirmed using epifluorescent microscopy. Spiked *Artemia* were then homogenized (PRO Scientific^®^, Series 250 Homogenizer) at max speed for 120 s, and homogenate was serial diluted (10-fold) in 1X PBS and spread plated using glass rattler beads (Zymo Rattler™ Plating Beads, 4.5 mm) onto thiosulfate citrate bile salts sucrose (TCBS) agar amended with 300 μg mL^−1^ kanamycin in duplicate. The addition of kanamycin to the TCBS plates selected against any non-GFP tagged *Vibrio* cells that may have been present. TCBS plates were incubated overnight at 30 °C. The resulting plate counts were used to calculate the approximate level of acquired dose.

### Uptake by *E. pallida*

To establish a connection between ingestion and *Vibrio* uptake, a controlled feeding study was conducted to measure the level of acquired GFP-tagged *Vibrio* spp. following exposure in a microcosm. Cultures of GFP-tagged *Vibrio* spp. and *Artemia* were prepared and combined as described above in *“Artemia* Colonization” to produce spiked *Artemia*. To increase the feeding opportunity, four *Artemia* spike exposures (~250 individuals each) were combined for a total exposure of ~1,000 individuals resulting in a maximum feeding dose (assuming ingestion of all *Artemia*) of ~1.96 x10^7^ CFU, ~5.89 × 10^6^ CFU, and 3.04 × 10^7^ CFU for *V. alginolyticus*, *V. harveyi*, and *V. mediterranei* trials, respectively (Table 2). Colonization of the spiked *Artemia* was confirmed prior to anemone feeding using fluorescent microscopy. Control *Artemia* (non-spiked) were prepared in tandem using the protocol but were inoculated with 50 μL of sterile 1X PBS instead of GFP-*Vibrio* spp.

**Table 2:** GFP-*Vibrio* spp. dosing patterns, *Artemia* acquisition efficacy, *Vibrio* exposure concentration, and recovered CFU from anemone homogenate. For each experimental trial ~1000 *Artemia* (individuals) and 6 anemones (individuals) were exposed.

| *Vibrio* spp. | Treatment Name | Total GFP-*Vibrio* spp. Exposed to *Artemia* (CFU)<sup>a</sup> | Mean GFP-*Vibrio* spp. Carried by *Artemia* (CFU/~1000 *Artemia*)<sup>b</sup> | Total GFP-*Vibrio* spp. Inoculated into Microcosm Water (CFU)<sup>a</sup> | Mean GFP-*Vibrio* spp. Recovered from *E. pallida* Homogenate (CFU mL<sup>-1</sup>) |
| --- | --- | --- | --- | --- | --- |
| *V. alginolyticus* | Spiked fed | 4.53 x 10<sup>8</sup> | 1.96 x 10<sup>7</sup> ± 5.60 x 10<sup>5</sup> | NA | 6.92 x 10<sup>4</sup> ± 7.48 x 10<sup>3</sup> |
| *V. alginolyticus* | Water exposed control fed | NA | NA | 4.53 x 10<sup>8</sup> | 3.33 x 10<sup>2</sup> ± 1.02 x 10<sup>2</sup> |
| *V. alginolyticus* | Water exposed not fed | NA | NA | 4.53 x 10<sup>8</sup> | 8.33 x 10<sup>1</sup> ± 4.00 x 10<sup>1</sup> |
| *V. alginolyticus* | Control | NA | NA | NA | 0.00 ± 0.00 |
| *V. harveyi* | Spiked fed | 2.75 x 10<sup>8</sup> | 5.89 x 10<sup>6</sup> ± 2.56 x 10<sup>5</sup> | NA | 2.59 x 10<sup>5</sup> ± 1.12 x 10<sup>5</sup> |
| *V. harveyi* | Water exposed control fed | NA | NA | 2.75 x 10<sup>8</sup> | 4.10 x 10<sup>3</sup> ± 1.61 x 10<sup>3</sup> |
| *V. harveyi* | Water exposed not fed | NA | NA | 2.75 x 10<sup>8</sup> | 8.33 x 10<sup>1</sup> ± 5.43 x 10<sup>1</sup> |
| *V. harveyi* | Control | NA | NA | NA | 0.00 ± 0.00 |
| *V. mediterranei* | Spiked fed | 6.05 x 10<sup>7</sup> | 3.04 x 10<sup>7</sup> ± 7.06 x 10<sup>5</sup> | NA | 1.67 x 10<sup>5</sup> ± 5.88 x 10<sup>4</sup> |
| *V. mediterranei* | Water exposed control fed | NA | NA | 6.05 x 10<sup>7</sup> | 5.90 x 10<sup>3</sup> ± 1.38 x 10<sup>3</sup> |
| *V. mediterranei* | Water exposed not fed | NA | NA | 6.05 x 10<sup>7</sup> | 3.83 x 10<sup>2</sup> ± 1.51 x 10<sup>2</sup> |
| *V. mediterranei* | Control | NA | NA | NA | 0.00 ± 0.00 |

<sup>a</sup>Total GFP-*Vibrio* exposure represents the CFU concentration introduced to *Artemia* to promote colonization this was calculated during *Artemia* dose assessment as the sum of four exposure trials (~250 *Artemia* each). Initial exposures were administered at 1.13 x 10<sup>8</sup>, 6.87 x 10<sup>7</sup>, and 1.51 x 10<sup>7</sup> CFU/~250 *Artemia* for *V. alginolyticus*, *V. harveyi*, and *V. mediterranei*, respectively. Water exposure trials were inoculated directly into the microcosm using the same concentration.
<sup>b</sup>Total GFP-*Vibrio* carried by *Artemia* represents the CFU concentration acquired by *Artemia* following exposure. This was calculated during *Artemia* dose assessment as the sum of four exposure trials (~250 *Artemia* each). Exposures resulted in an *Artemia*-acquired dose of 4.90 x 10<sup>6</sup>, 1.47 x 10<sup>6</sup>, and 7.59 x 10<sup>6</sup> CFU/~250 *Artemia* for *V. alginolyticus*, *V. harveyi*, and *V. mediterranei*, respectively.

Experimental microcosms were constructed to house the anemones during exposure trials. Microcosms were created using 18 × 12.5 × 5 cm Pyrex^®^ dishes filled with 750 mL of sterile ASW. Each microcosm contained a submerged six well tissue culture plate to provide substrate for *E. pallida* (N = 6 per treatment). Prior to exposure, experimental *E. pallida* were transferred to the microcosm chambers and allowed to acclimate for 18 h. Anemones used in experiments were selected based on size and consisted of individuals ranging from 1.5 cm to 3 cm (at full extension) to reduce the influence of feeding bias by large or small individuals. No discolored or wilting anemones were selected (see Figure S4 for an example of healthy *E. pallida* appearance). Care was taken during anemone detachment to ensure no damage to the tentacles or oral disc occurred. All anemones were checked visually for viability following acclimatization and replaced as needed. Experimental exposures were administered as detailed in Table 2 for a duration of 3 h. For trials were *Artemia* were fed to *E. pallida*, anemones were observed for the first 20 min following exposure to visually confirm ingestion. Anemones were checked every 30 min to ensure feeding behavior was continued and to stir microcosm water (to prevent *Artemia* from congregating out of anemone reach). Following exposure, anemones were collected from the chambers, transferred into individual 50 mL conical tubes containing 40 mL of sterile ASW, and vortexed for 30 s. This process was repeated twice to remove any non-ingested GFP-*Vibrio* cells. Washed anemones were then transferred into 10 mL of sterile ASW for homogenization. 100 μL of ASW was removed prior to homogenization and spread plated with glass rattler beads (Zymo Rattler™ Plating Beads, 4.5 mm) onto TCBS agar amended with 300 μg mL^−1^ kanamycin to ensure no ambient GFP-*Vibrio* (non-ingested) remained in the wash water (carry-over control). All anemones were then homogenized (PRO Scientific^®^, Series 250 Homogenizer) at max speed for 120 s. *E. pallida* homogenate was serial diluted (10-fold) in 1X PBS and spread plated with glass rattler beads onto TCBS agar amended with 300 μg mL^−1^ kanamycin, in duplicate. Plates were incubated at 30 °C for 18 h. The resulting plate counts (CFU/mL) were used to calculate the uptake of GFP-*Vibrio* cells by the anemones under each experimental condition. Culture results were summarized and visualized in Rstudio using the packages ‘tidyverse’ and ‘readxl.’ Feeding exposures were compared using a pairwise Wilcoxon rank-sum test with a Bonferroni correction for significance.

## Acknowledgements

We kindly thank Dr. Eric Stabb, for donating the helper and donor strains used to facilitate GFP tagging of the target *Vibrio* spp. We also acknowledge the work of Charlyn Shue, Rachel Phan, and Samantha Weatherly for their assistance with laboratory processing.

